# Aldehyde dehydrogenase 3A1 deficiency leads to mitochondrial dysfunction and impacts salivary gland stem cell self-renewal, differentiation and survival

**DOI:** 10.1101/2021.10.25.461994

**Authors:** Vignesh Viswanathan, Hongbin Cao, Julie Saiki, Dadi Jiang, Aaron Mattingly, Dhanya Nambiar, Joshua Bloomstein, Yang Li, Sizun Jiang, Manish Chamoli, Davud Sirjani, Michael Kaplan, F Christopher Holsinger, Rachel Liang, Rie Von Eyben, Haowen Jiang, Li Guan, Edward Lagory, Zhiping Feng, Garry Nolan, Jiangbin Ye, Nicholas Denko, Sarah Knox, Daria-Mochly Rosen, Quynh-Thu Le

**Affiliations:** Department of Radiation Oncology, Stanford School of Medicine, Stanford, CA 94305.; Department of Radiation Oncology, The University of Texas MD Anderson Cancer Center, Houston, TX 77030; Department of Cell and Tissue Biology, University of California San Francisco, CA 94143; Department of Microbiology & Immunology, Stanford University School of Medicine, Stanford, CA 94305; Buck Institute for Research on Aging, 8001 Redwood Blvd., Novato, CA 94945.; Department of Otolaryngology–Head and Neck Surgery, Stanford University School of Medicine, Stanford, CA 94305; Department of Chemical and Systems Biology, Stanford University School of Medicine, Stanford, CA 94305; The Ohio State University Wexner Medical Center and OSU Comprehensive Cancer Center, Columbus, OH 43210

## Abstract

Adult salivary stem/progenitor cells (SSPC) have an intrinsic property to self-renew in order to maintain tissue architecture and homeostasis. Adult salivary glands have been documented to harbor SSPC, which have been shown to play a vital role in the regeneration of the glandular structures post radiation damage. We have previously demonstrated that activation of aldehyde dehydrogenase 3A1 (ALDH3A1) after radiation reduced aldehyde accumulation in SSPC, leading to less apoptosis and improved salivary function. We subsequently found that sustained pharmacological ALDH3A1 activation is critical to enhance regeneration of murine submandibular gland after radiation damage. Further investigation shows that ALDH3A1 function is crucial for SSPC self-renewal and differentiation even in the absence of radiation stress. Salivary glands from *Aldh3a1*-null mice have fewer acinar structures than wildtype mice. ALDH3A1 deletion or pharmacological inhibition in SSPC leads to a decrease in mitochondrial DNA copy number, lower expression of mitochondrial specific genes and proteins, structural abnormalities, lower membrane potential, and reduced cellular respiration. Loss or inhibition of ALDH3A1 also elevates ROS levels and accumulation of ALDH3A1 substrate 4-hydroxynonenal (4-HNE, a lipid peroxidation product), leading to decreased survival of murine SSPC that can be rescued by treatment with 4-HNE specific carbonyl scavengers. Our data indicate that ALDH3A1 activity protects mitochondrial function and is important for the development and regeneration activity of SSPC. This knowledge will help to guide our translational strategy of applying ALDH3A1 activators in the clinic to prevent radiation-related hyposalivation in head and neck cancer patients.

## Introduction

Salivary glands function to produce and secrete saliva that aids in digestion of food and maintenance of oral health. In humans, there are three pairs of major salivary gland namely parotid, submandibular and sublingual that provide 90% of resting saliva volumes (1). Loss of function due to pathological atrophy of salivary glands (xerostomia) is a common complication associated with radiation therapy (RT) in head and neck cancer (HNC) patients. Xerostomia leads to difficulty in chewing, swallowing food, dental decay, weight loss and overall poor quality of life (2). Because of their proximity to the draining lymph nodes, the submandibular glands (SMGs) are most likely to be damaged by radiation, including intensity modulated radiotherapy (IMRT) (3,4). The only approved drug to prevent RT-related xerostomia is Amifostine, which is rarely used because of its significant side effects. Other interventions, such as pilocarpine for symptom relief, are minimally effective and require use for many years (5). This imminent therapeutic gap has led us to focus on understanding salivary gland biology in order to protect them from radiation-induced damage (6-9).

Morphologically, SMGs are primarily composed of ductal and acinar cells. Acinar cells can be mucous or serous types that produce saliva, which is then moved through an organized ductal system into the oral cavity via the Wharton duct. Label retaining experiments that mark slow cycling stem/progenitor cells lining the ducts have identified ductal stem cell markers, such as C-kit, Ascl3, Ck5 and Ck14 (10-12). Recently, the existence of self-renewing cells in the acinar compartment that could undergo self-duplication to regenerate and maintain homeostasis have also been described (13). Multiple reports investigating the response of salivary stem/progenitor cells (SSPC) to stressors using mouse salivary gland embryogenesis model, lineage tracing as well as adult murine salivary gland irradiation models have illustrated complex signaling mechanisms driving self-renewal, differentiation and regeneration (14).

We previously identified a novel role of aldehyde dehydrogenase 3A1 (ALDH3A1) in facilitating recovery of salivary gland function in irradiated murine SMG models (9). ALDH3A1 is one of the 19 isoforms of the ALDH family that is primarily known for its detoxification function in corneal epithelial cells (14). We demonstrated that increasing ALDH3A1 activity using its specific activator, Alda-341 (d-limonene), protected murine SMG SSPC from RT-induced toxic aldehydes, leading to less SSPC cell deaths and improvement of salivary gland function. Intriguingly, we noted that sustained ALDH3A1 activation with Alda-341 long after clearance of RT-induced toxic aldehydes is critical to enhance regeneration of murine SMG after RT damage. Since *Aldh3a1* is one of the top enriched genes found in SSPC compared to non-SSPC from SMGs (6,9), we hypothesize that it is also important for stem cell function and salivary gland development regardless of any stressor. Unexpectedly, we found that loss of *Aldh3a1* leads to mitochondrial dysfunction that can negatively affect self-renewal, differentiation and survival of murine SSPC.

## Results

### Enhanced ALDH3A1 activity after irradiation is essential for SSPC survival and regeneration

We have demonstrated that irradiation of murine SMGs reduced saliva function substantially, which could be partially rescued by treatment of ALDH3A1 activator, Alda-341, through a reduction of aldehyde load (9). If reduced aldehyde load from radiation is the main driver to mitigate RT damage and restore function, then short term treatment with the activator should suffice in sustaining improved saliva function. To address this hypothesis, we designed an experiment where Alda-341 treatment was started immediately after RT (30 Gy in 5 fractions), continued for 8 weeks, then either stopped and observed or continued for a total of 20 weeks (**Figure 1A**). We have previously reported that continued treatment for 20 weeks improved saliva production over the control irradiated group without drug treatment (9). Interestingly, early termination of Alda-341 at 8 weeks post RT resulted in a decline in saliva function at week 20 down to the level of the control group and much lower than that of the continous drug treated group (**Figure 1B**). These results suggest that apart from decreasing RT-induced aldehyde load, ALDH3A1 may play another role in the regeneration process, and sustained enzyme activation is a requisite to see maximum benefits. These findings in addition to our previous observation of high ALDH3A1 expression in SSPC lead to the hypothesis that ALDH3A1 plays a role SSPC function and survival.

**Figure 1:**
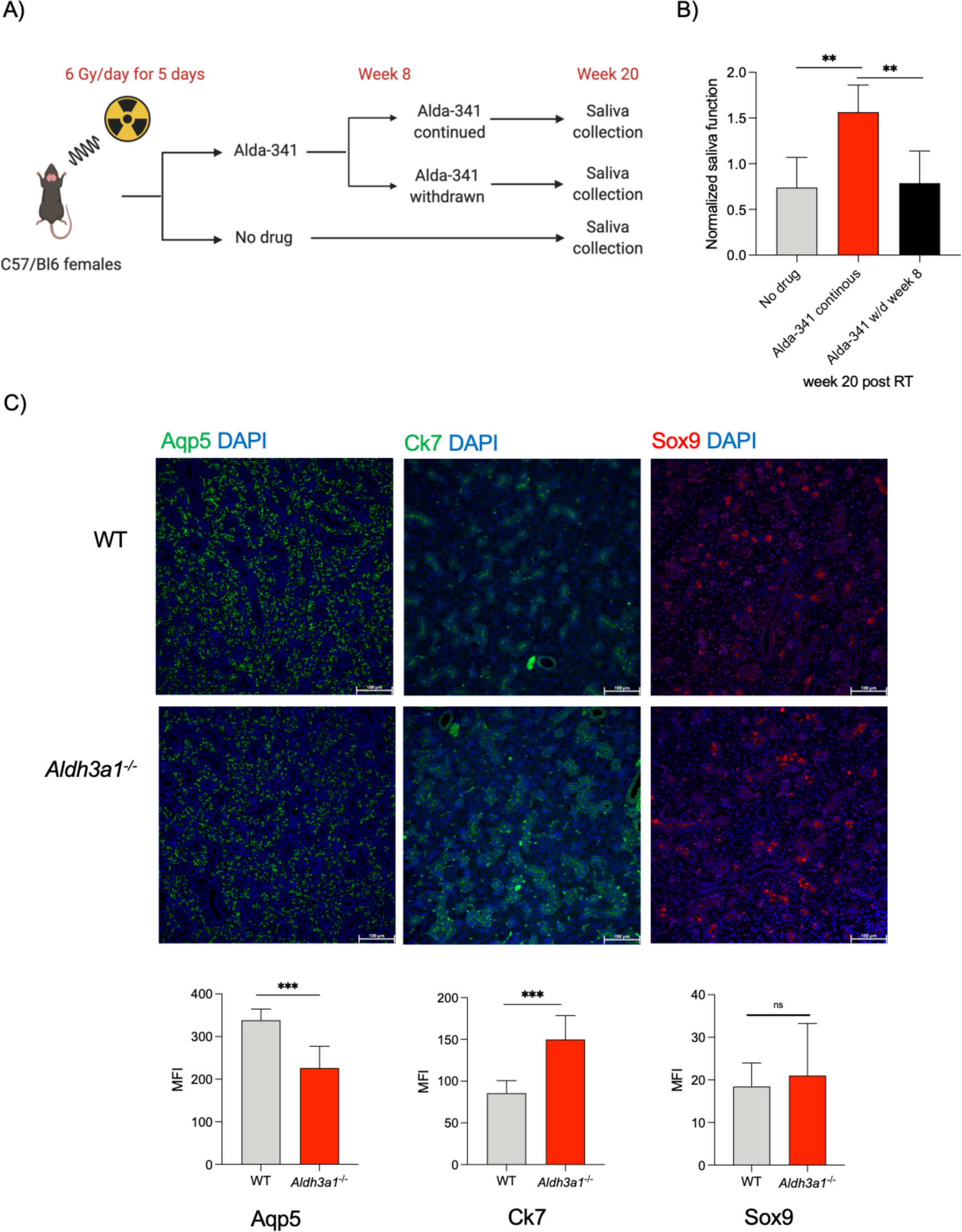
Sustained ALDH3A1 activity improves salivary function post RT in mice and its genetic deletion alters SMG tissue morphology. **A)** SMGs of mice were irradiated (30 Gy, 6 Gy x 5) followed by Alda-341 drug treatment for various time points upto 20 weeks. **B)** Normalized Saliva production in mice from different treatment groups at week 20 post 30 Gy radiation of salivary glands (6 Gy/day) (Alda-341 continous, Alda-341 withdrawn week 8, no drug control, N=7-8 per group). Student’s t-test was used to calculate the p value (** represents p value < 0.01). **C)** Immunofluorescent staining analyses of acinar (Aqp5) and ductal (Ck7, Sox9) markers in SMGs derived from WT and *Aldh3a1^-/-^* mice imaged at 200x total magnification that is quantified and represented as mean fluorescence intensity (MFI) in the lower panel (10 images per staining, n=3 mice/group). Scale bar: 100 μM. Student’s t-test was used to determine p value in panel **C**. Error bars represent SD. (*** represents p value < 0.001)

### *Aldh3a1* deficiency alters murine SMG tissue morphology in adults and branching morphogenesis of embryonic explants

We assessed differences in tissue morphology of the three major salivary glands: parotid, submandibular and sublingual in naïve *Aldh3a1^-/-^* mice as compared to WT mice. In both parotid and submandibular glands, we initially noted more ductal structures and possibly less acinar structures in the *Aldh3a1^-/-^* as compared to the WT mice on H&E staining (**Figure S1A)**. Using Ck7 and Sox9 as ductal markers and Aqp5 as acinar marker, we found that *Aldh3a1^-/-^* SMGs had fewer Aqp5^+^ acinar cells and more Ck7^+^ and Sox9^+^ ductal cells compared to WT SMGs (**Figure 1C**).

To investigate the role of *Aldh3a1* in development, we analyzed the expression pattern of *Aldh3a1* mRNA in murine SMGs at E14.5, 15.5, 16.5, P0 and P1 using RNAscope. Murine *Aldh3a1* expression was ubiquitious and diffuse in early stage (E14.5-15.5) and became limited and concentrated in the stem cell compartment marked by *c-kit* expression during later stages of development of murine SMGs (P0-1) as seen in **Figure2A**.We analyzed data from a published single cell RNA sequencing study done on murine embryonic and adult glands to identify the cell populations that expresses *Aldh3a1* during development, postnatal and adult murine SMG (15). UMAP plots show ubiquitous expression of *Aldh3a1* across different cell types at E12; the expression becomes restricted to specific cell populations at E14 and E16 (Krt19+ and basalduct+ cell cluster) (**Figure S2A**). In the postnatal and adult glands, Aldh3a1 was found to be enriched primarily in cell clusters of ductal origin (krt19+ duct, Basal duct and Gstt1+ ducts) (**Figure S2B**). The results bolster our findings that *Aldh3a1* expression is predominantly limited to ductal cells during development, postnatal and adult murine SMGs. Based on the morphological differences between *Aldh3a1^-/-^* and WT, we predicted that ALDH3A1 may influence murine embryonic SMG differentiation. SMGs isolated from E13.5 *Aldh3a1^-/-^* and WT mice were carefully dissected to remove the epithelium from the mesenchyme; the epithelial rudiments branch to form end buds when grown for 24 hours as described previously (11). We observed that *Aldh3a1^-/-^* murine embryonic glands had fewer end buds with associated smaller total epithelial area as compared to WT glands (**Figure 2B**). We also assessed the effect of a ALDH3A1 activator, Alda-341 (d-limonene), on branching ability of E13.5 murine SMGs and observed a significant dose-dependent increase in branching with more end buds in the Alda-341 treated group as compared to controls (**Figure 2C**). In the Alda-341 treated group, stem cell *c-kit* expression was found to be limited in the end buds as compared to the control (**Figure 2D**). Quantitative PCR analyses revealed that the phenotypic difference after Alda-341 treatment appears to be orchestrated by increased expression of genes regulating differentiation (*Aqp5, Etv5, Mist1*) and a decreased expression of stemness-related genes (*Sox2, Ck5*) (**Figure 2E**).

**Figure 2:**
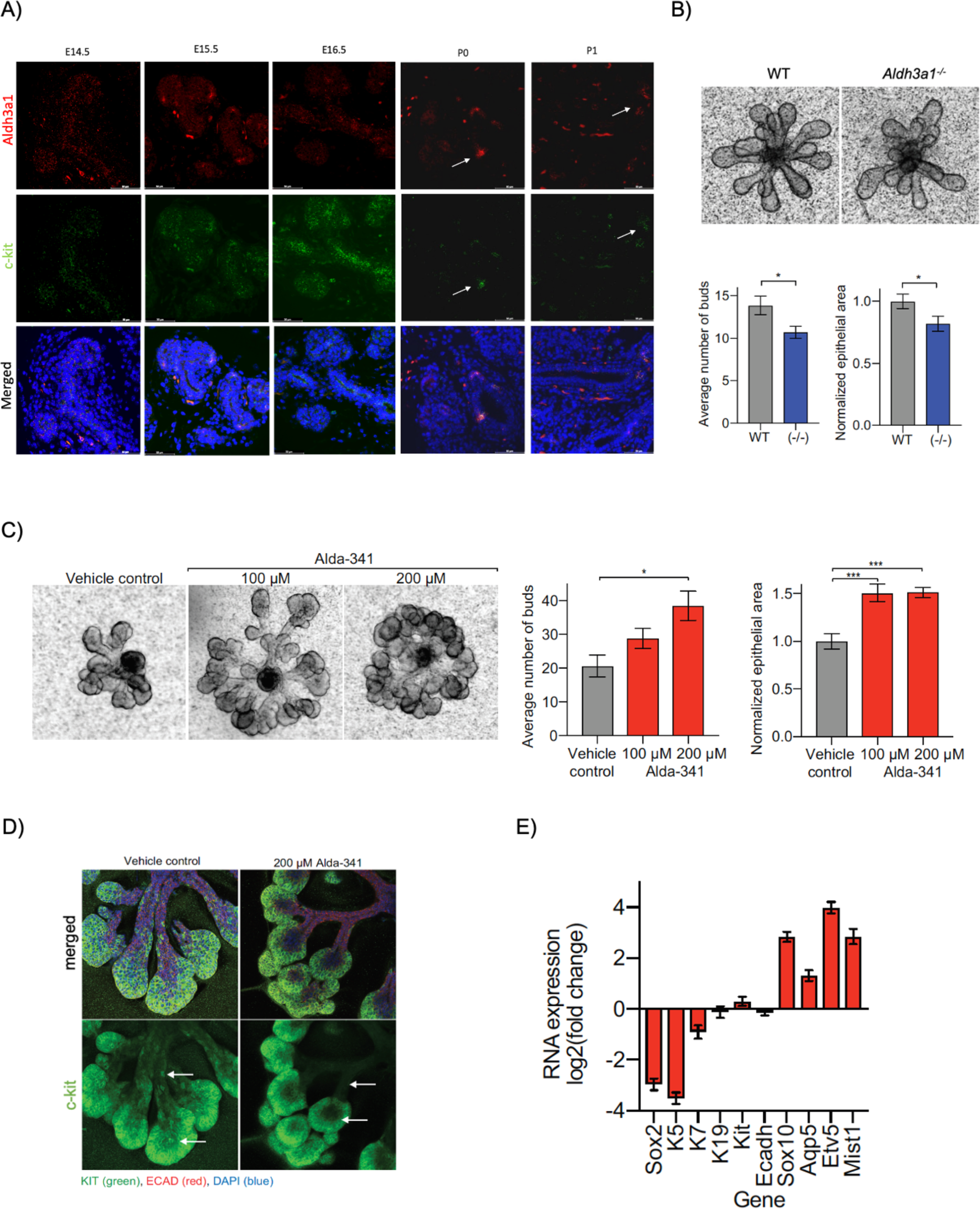
ALDH3A1 is expressed during development of embryonic salivary glands and is crucial for branching morphogenesis. **A)** Representative images of RNAscope hybridization analyses of ALDH3A1 (red) and c-kit (green) in embryonic salivary glands isolated at various stages of development. Arrow points at enrichment of ALDH3A1 and c-kit expression in salivary glands of postnatal (p0 and p1) mice. (5-6 epithelia/stage of development). Scale bar: 50 μM **B)** Upper panel represents embryonic SMG epithelia (E13.5) from WT (left) and *Aldh3a1^-/-^* (right) mouse embryos cultured for 24 h and imaged at 10x magnification. Lower left: Bud number counted from 17 WT and 16 *Aldh3a1^-/-^* epithelia. Lower right: Average epithelial area quantified from 15 WT and 14 *Aldh3a1^-/-^* epithelia using Image J (NIH) and normalized to WT. **C)** Representative images of E13.5 SMG epithelia from murine embryos were treated with vehicle control, 100 μM, and 200 μM Alda-341 (left to right), cultured for 24 h and imaged at 100x total magnification (N = 6-7 epithelia per group). Middle panel: Bud number was counted for each epithelium and averaged per group. Right panel: Epithelial area was quantified using Image J (NIH) and normalized to vehicle control. **D)** Representative images for vehicle control (left) and 200 μM Alda-341 (right) after 24 h in culture and immunoassayed with c-KIT (green), ECAD (red), and DAPI (blue), and imaged with a confocal microscope. Ten μM confocal section of all three markers (top) and averaged sections of c-KIT only (bottom)**. E)** Reverse transcription quantitative PCR of RNA extracted from 4 epithelia per group. RNA expression of epithelia treated with 200 μM Alda-341 represented as a log2 fold change over RNA expression of epithelia treated with vehicle control. Error bars represent SD. Student’s t-test was used to calculate the p value for panel **B** and **C** (* represents p value <0.05, ** < 0.01, *** < 0.001)

### ALDH3A1 activity is essential for self-renewal of murine and human SSPC

To determine if ALDH3A1 function is crucial for the growth properties of murine SSPC, we investigated the effect of ALDH3A1 deficiency in self-renewal *in vitro*. Primary dissociated cells from murine salivary gland derived from age and gender matched WT and *Aldh3a1^-/-^* mice were FACS sorted for cells with EpCAM/CD24 high expression, which has previously been shown to be enriched for SSPC (7). These cells were then grown on a layer of growth factor reduced (GFR) matrigel supplemented with stem cell growth media (**Figure 3A**). By day 10, *Aldh3a1^-/-^* SSPC gave rise to significantly fewer spheres as compared to WT, which was consistent across multiple passages (**Figure 3B**). To test the reverse, we treated murine SSPC with ALDH3A1 activator Alda-341. Treatment with Alda-341 significantly increased the number of spheres in a dose-dependent manner, as compared to the vehicle control-treated group (**Figure 3C**). This observation was also extended to human samples; treatment of SSPC from five patient-derived SMG cultures with two concentrations of Alda-341 similarly resulted in increased sphere formation compared to vehicle control in all five samples (**Figure 3D**).

**Figure 3:**
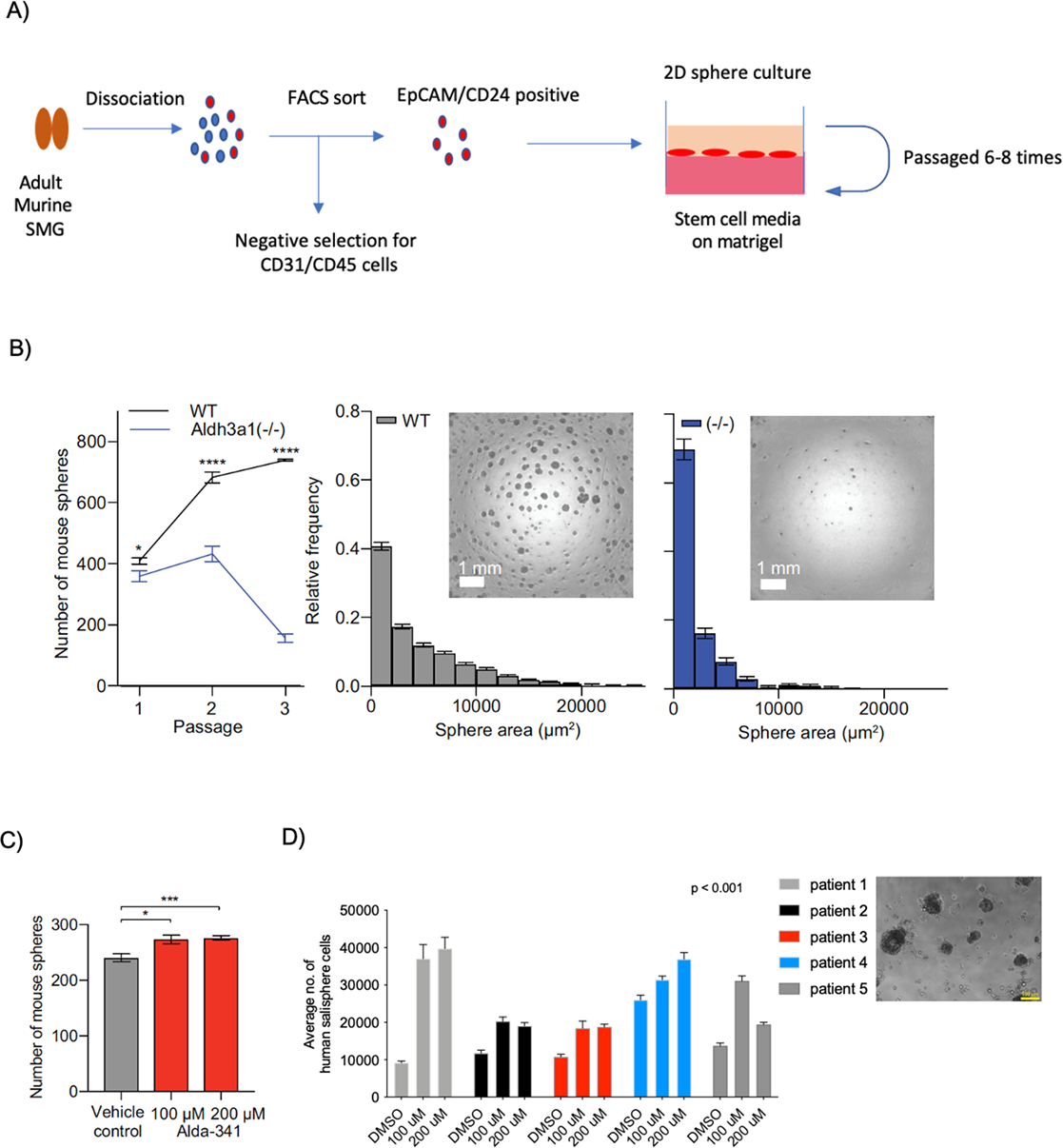
ALDH3A1 deficiency impacts self renewal and downregulates mitochondrial gene expression in SSPC. **A)** Schematic of SSPC isolation and culture from murine SMG. **B)** Left panel shows graph average number of spheres per well at day 7 was calculated by imaging each well and quantifying by Image J (NIH). Cells were passaged every 7 d for 3 passages (6 replicates per group). Middle and right panel represents graphs with average frequency over sphere area in WT and *Aldh3a1^-/-^ SSPC* with representative images of spheres from WT and *Aldh3a1^-/-^* SSPC **C)** Murine SSPC derived sphere count at day 7 post treatment with 100 μM or 200 μM Alda-341 or vehicle control. Sphere number per well quantified by Image J (NIH). (6 replicates per group). **D)** Human salivary gland derived SSPC count at day 7 in the presence of vehicle control, 100 or 200 μM Alda-341 (n=5 patients, 3 technical replicates). Representative image of human salivary gland spheres is shown. Error bars represent SD. Student’s t-test was used to calculate the p value (** represents p value < 0.01).

### Gene expression analyses identified dysregulated mitochondrial gene expression in *Aldh3a1^-/-^* SSPC

To identify molecular pathways impacted by *Aldh3a1* deficiency, we performed RNA-seq analyses on freshly isolated WT and *Aldh3a1^-/-^* murine SSPC (n=3/group). In total, 175 genes showed a significant log fold change with a FDR < 0.1 (**Figure 4A**). Gene Ontology analysis of these 175 genes using MetaCORE identified mitochondrial pathways to be significantly different in the *Aldh3a1^-/-^* SSPC as compared to WT (**Figure 4B**). Nine mitochondrial encoded genes were downregulated in the *Aldh3a1^-/-^* cells as compared to WT cells (**Figure 4C**). RNA-seq results of mitochondrial encoded genes were subsequently verified by qPCR using freshly isolated murine WT and *Aldh3a1^-/-^* SSPC (**Figure 4D**). Mitochondrial DNA copy number was lower in *Aldh3a1^-/-^* SSPC compared to WT, as reflected by the lower level of mitochondrial coded genes, *Cycs, Cox3* and *16rs,* normalized to nuclear coded gene, *Beta-globin* (**Figure S3A**). Immunostaining experiments showed reduced expression of mitochondrial markers Tom20 and VDAC1 in *Aldh3a1^-/-^* SMGs as compared to WT SMGs (**Figure 5A, 5B**). These differences in mitochondrial marker expression were also observed in primary spheres derived from *Aldh3a1^-/-^* and WT murine SSPC when analyzed by Immunostaining (**Figure S3B**). Ultrastructure analyses using transmission electron microscopy revealed reduced abundance of mitochondria and deformities in cristae and intramembranous space in of *Aldh3a1^-/-^* SMGs (**Figure 5C**). We also observed more mitochondria undergoing mitophagy in *Aldh3a1^-/-^* SMGs using TEM (**Figure 5D**). Mitophagy marker, Parkin1, was increased in *Aldh3a1^-/-^* SMGs as compared to WT, suggesting increased mitophagy (**Figure 5E**). These results validate our RNA-seq analyses and suggest a novel role of the cytosolic ALDH3A1 in influencing mitochondrial content, structure, protein expression and dynamics in murine salivary glands.

**Figure 4:**
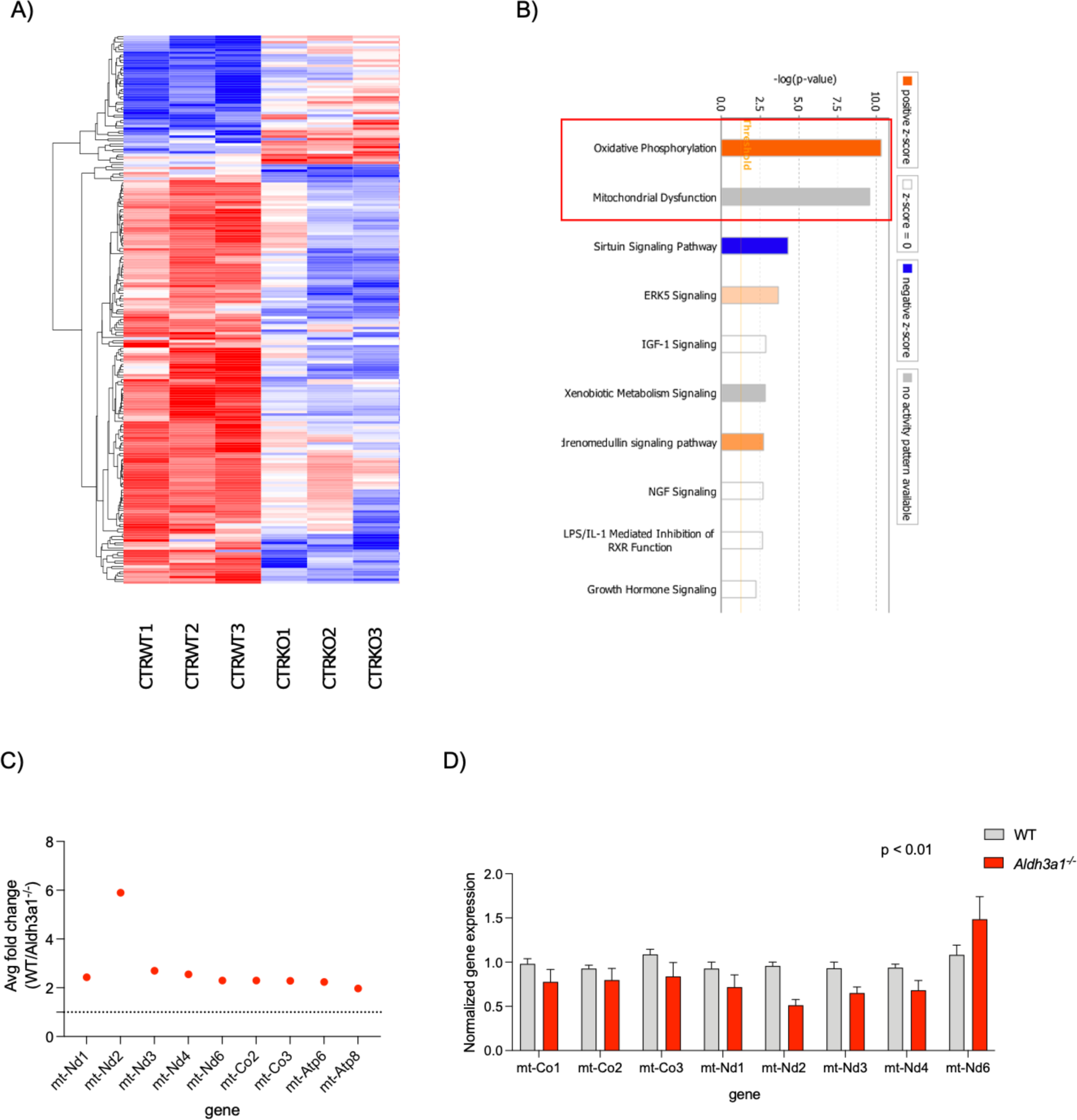
**A)** Heatmap showing differences in gene expression profile of WT and *Aldh3a1^-/-^* sorted salivary stem cells (n=3/group). **B)** Gene Ontology MetaCORE analyses of top 175 differentially regulated genes (FDR < 0.1) showing enriched differential pathways between WT and *Aldh3a1^-/-^* SSPC. C) Average fold change in expression of mitochondrial genes in WT as compared to *Aldh3a1^-/-^* SSPC. **D)** Quantitative PCR analyses of mitochondrial genes in murine SSPC derived from WT and *Aldh3a1^-/-^* SMGs (n=3 mice per group).

**Figure 5:**
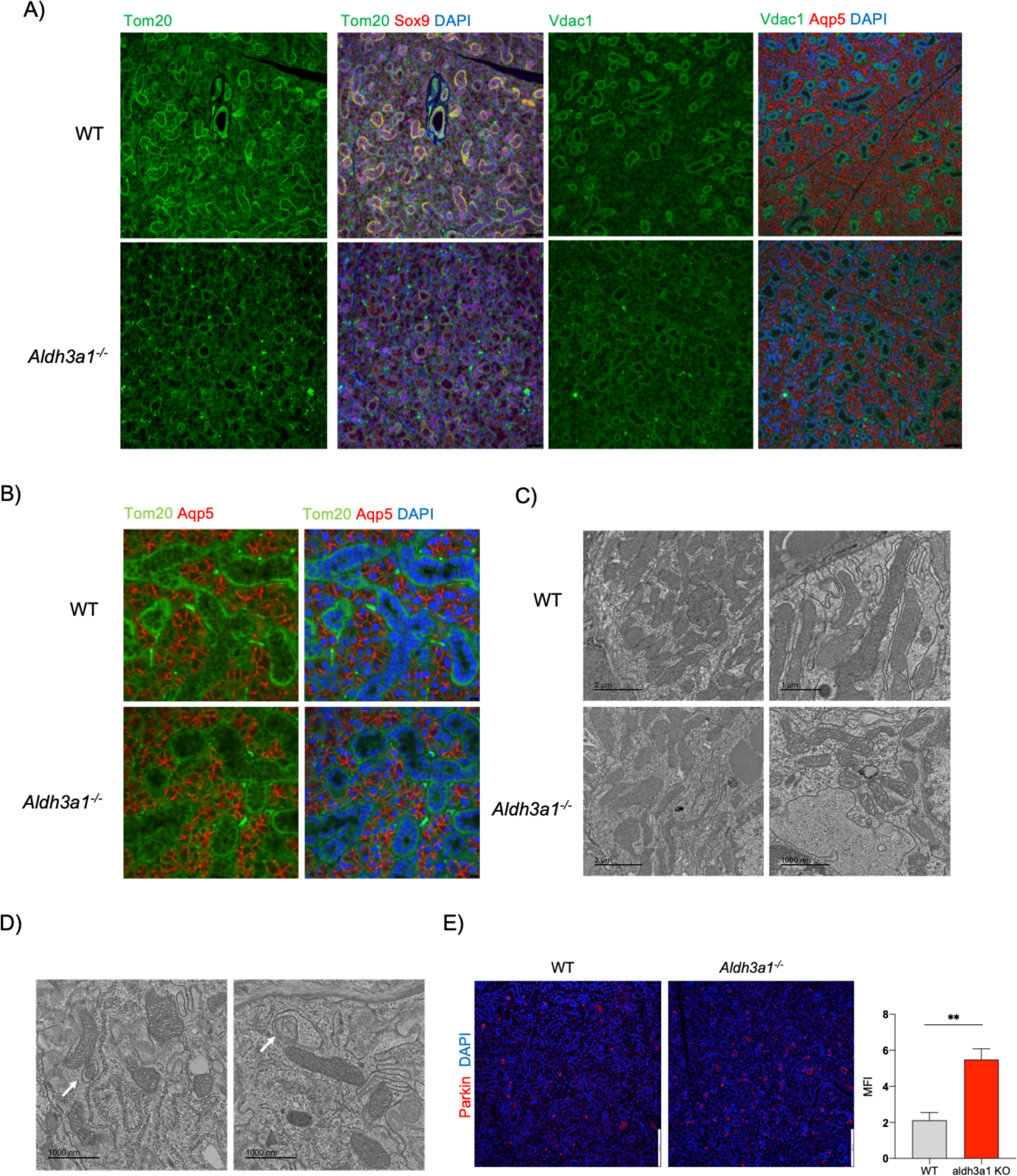
**A)** Representative Immunostaining images of mitochondrial (Tom20, Vdac1), ductal (Sox9) and acinar (Aqp5) markers in WT and *Aldh3a1^-/-^* SMGs. Scale bar: 50 μM (10 images per staining, n=3/group). **B)** High magnification immunostaining image of WT and *Aldh3a1^-/-^* SMGs showing mitochondrial marker Tom20 and Acinar marker Aqp5. **C)** TEM image of SMGs derived from WT and *Aldh3a1^-/-^* showing mitochondrial abundance and ultrastructure (n=3/group, scale bar= 2 μM left panel, 1 μM right panel). D) Representative TEM image of *Aldh3a1^-/-^* SMG showing mitophagy. E) Immunostaining analyses of mitophagy marker, Parkin, in WT and *Aldh3a1^-/-^* SMGs imaged at 200x total magnification that is quantified and represented in the right panel (n=3/group, scale bar: 130 uM). Error bars represent SD. Student’s t-test was used to calculate the p value (** represents p value < 0.01).

### ALDH3A1 impacts mitochondrial function of murine salivary gland cells

We assessed alterations in mitochondrial functional attributes in terms of membrane potential, sensitivity to mitochondrial inhibitor and respiration in murine salivary cells with decreased or no ALDH3A1 activity in comparison to controls. Loss of membrane potential in mitochondria is linked with mitochondrial dysfunction. Using the JC-1 assay, we found that *Aldh3a1^-/-^* SSPC had a significantly lower average membrane potential as compared to WT SSPC (**Figure 6A**). *Aldh3a1^-/-^* SSPC were also two folds more sensitive to a mitochondrial uncoupler, FCCP, as represented by a significantly higher percentage of positive cells undergoing apoptosis at 24 hours after treatment as compared to WT SSPC (**Figure 6B**). Differences in mitochondrial integrity and protein expression could translate to deficiency in mitochondrial primary function, which is cellular respiration and ATP production. We could not employ our sphere model for this experiment due to constraints of growth requirements and adherent conditions. To overcome this, we used a normal murine salivary gland cell line (mSGc) as an alternate model. mSGc can be maintained for multiple passages without a loss of proliferation potential, readily form 3D-spheroids and importantly express a panel of well-established salivary gland epithelial cell markers (16). We used a previously reported specific inhibitor of ALDH3A1 activity, CB29 (17), to mimic the *Aldh3a1^-/-^* phenotype in the cell line model. The EC50 of the inhibitory effect of CB29 to ALDH3A1 was determined by a dose-response study (**Figure S4A, B**). To test the effect of ALDH3A1 inhibition on mitochondrial respiration, we treated mSGc cells with CB29 and measured oxygen consumption rate (OCR) to estimate basal, maximum and ATP production using a Seahorse XF analyzer 24 hours post treatment. We found a significant decrease in ATP production as well as basal and maximum respiration in the CB29 treated cells as compared to vehicle control. Also, treatment with ALDH3A1 activator, Alda-341, alone increased the basal and maximum respiration, and when given in combination with CB29 – partially rescued the effect of the inhibitor on mitochondrial respiration and ATP production (**Figure 6C,D** **and E)**. CB29 treatment of WT SSPC also led to more apoptosis with associated decreased sphere formation as compared to vehicle control (**Figure S4C, D**). Complementaing the experiments above, we also used siRNA approach against *Aldh3a1* to confirm that reduced *Aldh3a1* expression lead to increased apoptosis and decreased sphere formation efficiency in mSGc cells (**Figure S4E, F and D).**

**Figure 6:**
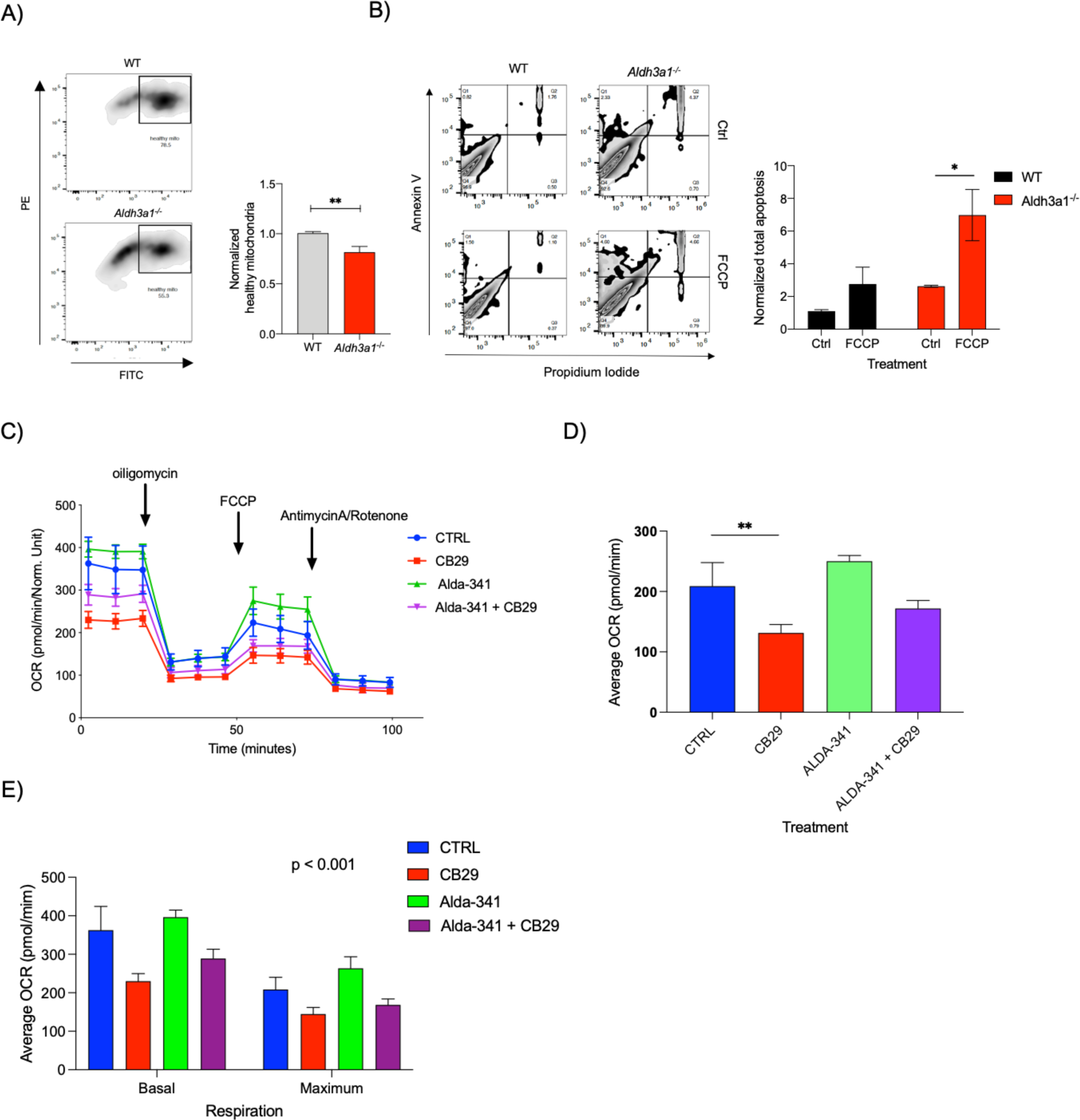
**A)** JC-1 assay analyses of WT and *Aldh3a1^-/-^* murine SSPC showing proportions of PE-high/FITC-high cells having mitochondria with intact membrane potential that is quantified and represented in the graph in right panel (n=3, 2 technical replicates). **B)** Quadrant plots showing Annexin-V/PI staining of WT and *Aldh3a1^-/-^* murine SSPC 24 hours post treatment with mitochondrial inhibitor FCCP, which is quantified and represented in Graph in right panel (n=3 with technical duplicates)**. C)** Average OCR over time of mSGc cells treated with vehicle, 10uM ALDH3A1 inhibitor CB29, 100uM Alda-341 and combination of both. **D)** Represents average ATP production in various treatment groups. (n=2 with 4 technical replicates). **E)** represents average Basal and Maximum respiration in various treatment groups. Error bars represent SD. Student’s t-test was used to calculate the p value for panel **A, B and D**. One way ANOVA was used to determine p value for panel **C**. (*represents p value < 0.05, ** < 0.01, *** < 0.001)

### Levels of ROS and substrate 4-HNE influences survival of *Aldh3a1^-/-^* SSPC

Increased ROS levels have been associated with mitochondrial dysfunction. In the murine SSPC model, ROS levels were significantly higher in the *Aldh3a1^-/-^* SSPC as compared to WT SSPC (**Figure 7A**). We have previously found that *Aldh3a1^-/-^* SSPC have higher basal apoptosis as compared to WT cells (9). To demonstrate that ROS accumulation impacts survival, we tested the effect of antioxidant treatment on survival of murine *Aldh3a1^-/-^* SSPC. Treatment of *Aldh3a1^-/-^* cells for 24 hours with mito-TEMPO, a mitochondria-specific ROS scavenging molecule, significantly improved *Aldh3a1^-/-^* SSPC survival as compared to vehicle control (**Figure 7B**). Glutathione plays anti-oxidative role against elevated ROS. ALDH3A1 helps in converting oxidized glutathione (GSSG) to reduced form (GSH) (18). As expected, in the absence of ALDH3A1, we observed a decreased glutathione turnover in murine *Aldh3a1^-/-^* SSPC by LC/MS analyses (**Figure 7C**), which can contribute to oxidative stress. One of the main substrates of ALDH3A1 is 4-hydoxynonenal (4-HNE) which is a product of lipid peroxidation (LPO). We observed increased accumulation of 4-HNE in SMGs of *Aldh3a1^-/-^* as compared to WT glands as shown by Immunostaining and Western blotting (**Figure 7D, E**). Hydrazine derivatives have been reported to rescue the effects of 4-HNE accumulation in smooth muscle cells (19). As expected, hydralazine treatment on *Aldh3a1^-/-^* SSPC for 24 hours significantly decreased the proportion of cells undergoing apoptosis and improved survival (**Figure 7F**). To confirm if the hydrazine acts like a scavenger for 4-HNE, we exposed WT SSPC for 24 hours with 4-HNE alone, hydralazine alone, or the two together and assessed apoptosis. There was an induction of cell death with 4-HNE treatment alone, which was rescued upon co-treatment of hydralazine (**Figure 7G**).

**Figure 7:**
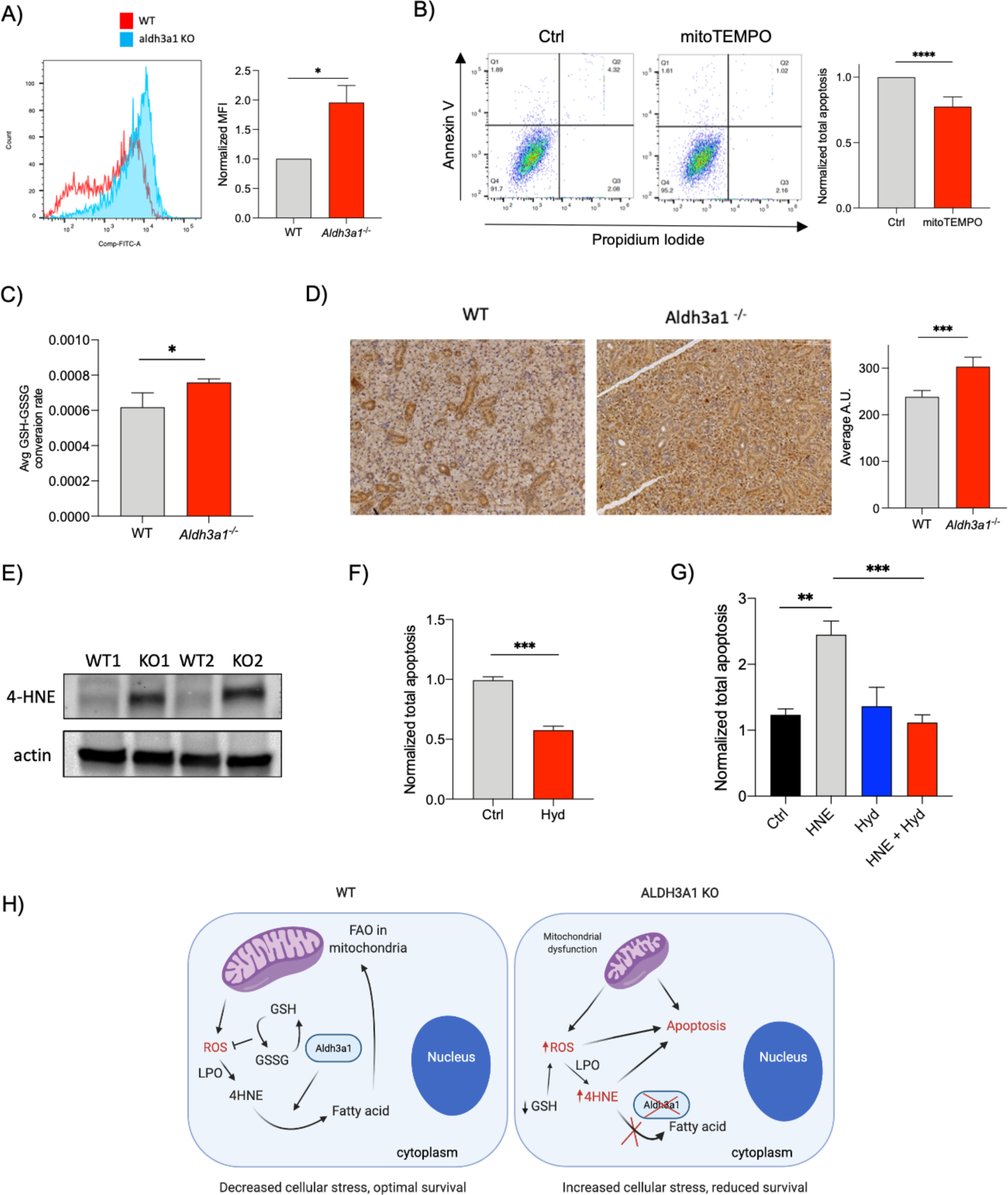
**A)** Histogram plots of ROS levels determined by FACS analyses in WT and *Aldh3a1^-/-^* murine SSPC that is quantified and represented in the right panel (n=3, 2 technical replicates). **B)** Bar graph showing normalized percent apoptosis of *Aldh3a1^-/-^* murine SSPC 24 hours post treatment with vehicle control and 100nM mito-TEMPO. (n=2, 3 technical replicates). **C)** Carbon labelling experiments demonstrate differences in glutathione turnover in WT and *Aldh3a1^-/-^* SSPC. **D)** Immunostaining and **(E)** Western blot analyses of 4-HNE in WT and *Aldh3a1^-/-^* SMGs (n=3/group for IHC, n=2 for Western blotting). Scale bar: 100 μM **E)** Average percent apoptosis in *Aldh3a1^-/-^* SSPC 24 hours post treatment with 10 uM of Hydralazine normalized to control. (n=3, 3 technical replicates). **F)** Average percent apoptosis in WT SSPC 24 hours post treatment with 10 uM HNE, 10 uM of Hydralazine and combination with HNE and Hydralazine normalized to control (n=2, 3 technical replicates). Error bars represent SD. Student’s t-test was used to calculate the p value for panel **A, B, C, D and F**. One way ANOVA was used to determine p value for panel **G**. (*represents p value < 0.05, ** < 0.01, *** < 0.001). **H)** Illustration representing potential role of ALDH3A1 in regulating mitochondrial function and survival of murine SSPC. ROS a derivative of mitochondrial activity, when accumulated can mediate lipid peroxidation (LPO) to make 4-HNE. 4-HNE, one of the major substrates of ALDH3A1 is broken down to fatty acids. ALDH3A1 converts GSSG to GSH which can act as an antioxidant to reduce oxidative stress. In the absence of ALDH3A1, reduction of GSH reserve, accumulation of ROS and its by-products 4-HNE can impair mitochondrial function to reduce overall survival of murine SSPC.

## Discussion

Gene expression profiling of murine SSPC helped us to identify and validate GDNF and ALDH3A1 as crucial determinants of salivary gland function and repair upon radiation stress (7,8). ALDH3A1-deficient mice display poorer saliva function after RT as compared to WT, and treatment with the ALDH3A1 activator, Alda-341 (D-limonene), improved regeneration of SMGs by reducing aldehyde load and promoting survival (9). In this study, we found that ALDH3A1 is important for self-renewal, survival and differentiation of murine salivary gland stem/progenitor cells without radiation stress. Genetic deletion of *Aldh3a1* results in morphological differences in murine SMG that can be attributed to abnormal development patterns. Branching morphogenesis of murine embryonic SMGs is an excellent model to study the effect of cell specific ablation of factors important for differentiation or development (12). Reduced branching and number of end buds in *Aldh3a1* deficient explants suggests its role in acinar cell development.

These findings resonate with other studies; for example, SOX2 was identified as a marker of SMG progenitor cells that are capable of giving rise to acinar cells but not ductal cells (20). Indeed, differences observed in embryonic morphogenesis of fetal explants can explain why we see reduced acinar compartment in adult SMGs of KO mice as compared to WT. ALDH3A1 activity also promoted self-renewal of murine and human SSPC. Self-renewal is a characteristic attribute of adult stem cells, which upon division gives rise to a differentiated daughter cell and a stem cell. Sphere formation assay from isolated murine SSPC is an *in vitro* technique to assess changes in self-renewal upon treatment or intervention (21). As sphere formation is a balance of proliferation and survival, we predict that ALDH3A1 deficiency affects SSPC survival, thus shifting the overall balance and reducing sphere formation.

Unexpectedly, in-depth gene expression analyses and functional studies identified strong link between *Aldh3a1* deficiency and reduced mitochondrial gene expression and function in SSPC. A survey of the literature showed only two studies that reported a relationship between ALDH3A1 and mitochondrial function. In a yeast model, a homolog of ALDH3A1 plays a critical role in the synthesis of a precursor molecule of coenzyme Q, an important carrier of the mitochondrial electron transport chain (ETC) (22). In gastric cancer, inhibiting ALDH3A1 impairs mitochondrial activity due to reduced beta oxidation of lipids and acetyl co-A flux for TCA cycle (23), which was also demonstrated by our LC/MS analyses (**Figure S3C**). Moreover, increased ALDH3A1 activity by Alda-341 (d-limonene) treatment in mice upregulated mitochondrial related genes in WT SSPC as demonstrated by our independent gene expression analyses (**Figure S3D and E**). These cumulative findings strongly demonstrate that activation of the cytosolic ALDH3A1 is required and sufficient to support normal mitochondrial function in salivary stem/progenitor cells under basal, non-stressed conditions.

Mechanistically, our data suggest a vicious cycle between loss of ALDH3A1 activity and mitochondrial function: increased ROS levels in *Aldh3a1^-/-^* SSPC leads to higher oxidative stress, reduced glutathione turnover, increased HNE accumulation and mitochondrial damage and reduced survival (**Figure 7H**). Mitochondria are the primary source of ROS, which is partially converted to toxic products such as 4-Hydroxynonenal (4-HNE) via lipid peroxidation (LPO) (24,25). ALDH3A1 inactivates the ROS product, 4-HNE, as well as replenishes reduced glutathione pool to maintain homeostasis and viability (18, 26). the loss or decrease activity of ALDH3A1 leads to accumulation of 4-HNE, which in turn causes protein adduct formation and inactivation, mitochondrial dysfunction, increased ROS accumulation and increased apoptosis in cells (26,27).

Our data showed that early termination of Alda-341 treatment at 8 weeks after radiation led to declining salivary function whereas, continued treatment for 20 weeks results in maintenance of improved saliva function. This can be an attributed to the role of ALDH3A1 acitivy in stem cell renewal after a physiologic insult such as radiation. Physiologically, stem cells are found in hypoxic niche thus normally these cells experience low ROS levels (28). However, upon radiation, increased ROS levels impairs their ability to self-renew and proliferate (29,30). We have demonstrated that improved ALDH3A1 activity can protect against ROS, therefore is even more critical for stem cell survival during this stage. Future work will explore how ALDH3A1 activity influences longterm effects such as chronic inflammation post radiation that poses a greater risk for tissue regeneration (31). Overall, our data helped to guide the duration of d-limonene treatment in a phase I trial that is about to be iniitated in head and neck cancer patients, in which d-limonene is to be administered during and after definitive chemoradiotherapy for salivary gland protection from radiation damage.

In summary, our data indicate that ALDH3A1 activity is required to support differentiation and function of salivary stem cells, at least in part, by maintaining mitochondrial number and functions. It explains for the differences in tissue architecture when this enzyme is deleted and supports the continuing use of ALDH3A1 activators after the end of radiation to assist stem cell renewal and differentiation in that setting.

## Methods

### Animals used

C57BL/6 mice (7–10 weeks) were purchased from Jackson Laboratories (Bar Harbor, ME). Experiments were done with 8-12 weeks old female mice. All the protocols were approved by The Administrative Panel on Laboratory Animal Care (APLAC) at Stanford University, Stanford, CA. All the experiments involving animals were done in adherence to the NIH Guide for the Care of and Use of Laboratory Animals. For our analyses, we used female mice because murine SMGs display sexual dimorphism and female SMGs show closer resemblance to humans (32).

### Salivary gland isolation and culture from mouse and human SMGs

Murine salivary glands were dissected and isolated from euthanized mice as described previously (8). Please see Supplemental methods for details.

### Immunostaining

Sections from paraffin embedded salivary gland tissue or spheres were used for Immunostaining. Slides were de-paraffinized by 2 rounds of incubation in fresh xylene for 10 min each. The slides were sequentially rehydrated by washing them in increasing concentration of water in alcohol for 3 min each. Antigen retrieval was performed by boiling the slides in 1x Antigen retrieval solution (Vector labs, Burlingame, CA) using rice cooker for 12 minutes. After cool down at room temperature slides were washed twice with PBS. For intracellular staining, slides were incubated in 0.1% Tween-20 in PBS (PBS-T) for 10 min and washed twice again with PBS. For immunohistochemistry (IHC), an addition step of treatment of slides with 3% hydrogen peroxide in methanol for 10 min was added followed by PBS wash. Slides were incubated blocking buffer for 1 hour at RT followed by overnight incubation at 4 C in the primary antibodies. List of antibodies and dilutions are provided in **Supplementary Table 1**. Next day, Slides were washed in PBS three times and then incubated in secondary antibody for 1 hr. Following secondary antibody incubation, slides were washed three times in PBS and Gold-anti FADE mounting media containing DAPI was added as a counterstain for immunofluorescence (IF). For IHC, slides for incubated in activated DAB for 30-60 sec and followed by multiple washes with distilled water. Slides were then de-hydrated by treating with decreasing amount of water in alcohol and then to xylene for 30 seconds each. Images were taken from ten random field of view using the Leica Dmi8 fluorescence microscope. These images were used for quantifications using ImageJ software (NIH, Bethesda, MD).

### Western Blotting

Protein was isolated from cells using 1X RIPA lysis buffer (EMD Millipore, Burlington, MA) containing 1X protease inhibitor cocktail (Thermo-Fisher, Waltham, MA). Protein concentrations were estimated using Pierce BCA assay kit (Thermo-Fisher). Thirty micrograms of protein were loaded in Mini-Protean precast gels (Bio-Rad, Hercules, CA). Gel electrophoresis was run at a steady voltage of 75 V at 4 C. After the gel run was completed, resolved proteins were transferred on a nitrocellulose membrane using the Turbo Transblot kit (Bio-Rad) following the manufacturer’s protocol. The blots were incubated in 3%BSA in TBS-T for 1 hours for blocking followed by overnight incubation in primary antibody. Next day, the blots were washed thrice in 0.1% Tween-20 in Tris Buffed Saline (TBS-T) for 5 minutes each. HRP conjugated secondary antibody was prepared in TBS-T (1:1000) added to the blots for incubation for 1 hour. Excess unbound secondary antibody was removed by three washes of TBS-T. Blots were developed by using Super Signal West Pico plus Chemiluminescent substrate (Thermo-Fisher).

### PI/Annexin V assay

Cells were washed with cold PBS and incubated with anti-Annexin V-FITC diluted solution (Bio-legend, San-diego, CA) for 10 minutes at 37 C in the dark followed by PI solutions for 5 min. Cells were washed again with cold PBS and subjected to FACS analyses using BD LSR FortessaX-20 flow cytometer. Using FACS DIVA software, positive quadrant gates in the plots were determined using unstained cells as a control sample. Cell populations were identified as early apoptosis (Q1:Annexin V ^+ve^/ PI^-ve^), late apoptosis (Q2:Annexin V^+ve^/ PI^+ve^) and total apoptosis as the sum of all three quadrants (Q1,Q2,Q3). Changes in apoptosis were analyzed after 24-48 hours of treatment with drugs (FCCP, CB29, mito-TEMPO, 4-HNE, Hydralazine) or vehicle control. FlowJo (Ashland, OR) was used for analyses and data was represented as average percent apoptosis in the treatment normalized to control.

### RNA Sequencing

RNA Sequencing experiment was done as previously described (9). Please see **Supplemental methods** for details.

### Single-Cell RNA-Seq data analyses

Ready to use SEURAT objects for Embryonic and Postnatal SMG intergrated datasets were retrieved from figshare as indicated in the original study (15). The code used for the analyses is provided in **Data S1**.

### RT-PCR

RNA was isolated from whole tissue or SSPC using RNAqueous Micro Kit (Ambion, Austin, TX). Total RNA samples were DNase-treated (Ambion) before complementary DNA (cDNA) synthesis using SuperScript reagents according to manufacturer’s protocol (Invitrogen, Waltham, MA). SYBRgreen RT-qPCR was performed using cDNA and primers used before (33) or designed using Primer3 and Beacon Designer software or found using PrimerBank (http://pga.mgh.harvard.edu/primerbank/). Gene expression was normalized to the housekeeping gene S29 (Rps29) or actin.

### Embryo salivary gland dissection and branching morphogenesis assay

Salivary glands were dissected out during various stages of murine embryos development as described previously (34). E13.5 salivary glands were used for the branching morphogenesis assay. Please refer to **Supplemental methods** for more details.

### EM imaging

Salivary glands from WT and *Aldh3a1^-/-^* mice were dissected and used for transmission electron microscopy. Please refer to **Supplemental methods** for details.

### Mitochondria respiration

Ten thousand cells were seeded in Agilent (Santa Clara, CA) Seahorse XF96 cell culture microplates. Next day, cells were treated with vehicle control, CB29 (20 μM), Alda-341 (100 μM), and combination of the two drugs. After 24 hours of Incubation, media was replaced, and cells were maintained at 37 C. Extracellular flux assay to measure mitochondrial respiration and ATP production was carried out as recommended by manufacturer’s protocol (Agilent). SRB assay (35) was used to quantify cell concentration in samples for normalization of data.

### Mitochondrial isolation and copy number analyses

Mitochondrial DNA was isolated from a modified version of a protocol (36). Cells were scraped using RIPA lysis buffer and incubated with 5 μl of Proteinase K (Invitrogen) for 55 C for 3 hours. Samples were sonicated briefly to shear the DNA. The debris was removed from spinning down the samples at 8000 g for 15 min. To the supernatant, equal volume of phenol/chloroform/Isoamyl alcohol mixture (25:4:1) was added. The samples were mixed well by vigorous shaking followed by centrifugation at 8000 g for 15 min. The clear upper layer was collected, added to equal volume of chloroform/isoamyl alcohol and mixed well. The samples were spun down and the upper layer was transferred to a fresh tube, to which 40 μl of 3M sodium acetate and 440 μl of Isopropanol were added. Samples were incubated at -20 C for 10 min to facilitate DNA precipitation. Samples were spun again to pellet the precipitated DNA. Supernatant was discarded and the pellet was washed with 70% alcohol. The supernatant was discarded, and the dried pellet was re-suspended in molecular grade water (Invitrogen). For copy number analyses, 1 ng of DNA was diluted in 100 μl of water. MtDNA and genomic DNA isolated was subjected to qPCR (standard conditions) using primers against mitochondrial genes cycs, cox3 and 16rs and nuclear encoded gene beta-globin. Primer sequences and run conditions for mitochondrial genes were as previously described (36). The final data was represented as relative amount of mitochondrial gene amplification normalized to nuclear gene amplification.

### Mitochondria membrane potential assay

JC-1 assay (Adipogen life sciences, San Diego, CA) was used to assess mitochondrial membrane potential. SSPC were incubated with 0.5 μM of JC-1 reagent in 3% Bovine Serum Albumin (BSA) in PBS for 10 min at 37 C. Cells were washed once with excess PBS and were resuspended in 200 μl of BSA solution. Samples were subjected to FACS analyses where dot plots with axes FITC-A vs PE-A with positive gates set according to the unstained sample. Using FlowJo software (BD biosciences, San Jose, CA) for analyses, healthy mitochondria were identified as cell populations that were FITC ^high^ PE ^high^ and unhealthy mitochondria (due to loss of membrane potential) were identified as FITC ^high^ PE ^low^ population.

### Estimation of ROS levels

Cellular ROS was estimated using the CM-H2DCFDA reagent (ThermoFisher). Cells were incubated with 5 μM of the reagent in 3%BSA in PBS at 37 C for 30 minutes. Cells were washed once with ice-cold PBS and then subjected to FACS analyses. Histogram plots of FITC-Area from 10,000 cells were recorded and mean fluorescence intensity (MFI) were estimated for analyses using FlowJo software.

### RNAscope RNA hybridization

Paraffin embedded tissue sections were used for RNAscope. Probes against mouse *Aldh3a1* and *ckit* were used (ACD bio, Newark, CA). The slides were baked at 70 C for an hour. Slides were deparaffinized with fresh xylene treatment followed by alcohol and finally rehydrated in distilled water. Antigen retrieval step was performed by incubating the slides in 1X RNAscope antigen retrieval buffer at 97 C for 30 min. After cool down, diluted protease reagent were added on to the sections and incubated at 40 C for ten min. Sides were washed with distilled water two times for 2 min each. Slides were incubated in the probes (1:50) at 40 C for 16 hours (overnight step). Next day, slides were washed twice in 1X RNAscope wash buffer 2 min each. Three step amplification was carried for 15-30 minutes at 40 C with intermittent washing steps. After the final wash, slides were incubated in washing buffer containing DAPI (1:10000) followed by mounting cover slips carefully with ProlongGold on glass slides. Five to Six epithelia images were acquired using a total magnification of 630X using the Leica DMi8 fluorescence inverted microscope.

### ALDH activity assay

Human recombinant ALDH1A1, ALDH1A2, ALDH1A3, ALDH1B1, ALDH2, ALDH3A1, ALDH3A2 and ALDH4A1 were expressed and purified using nickel column chromatography as previously described (37). The assay buffer was 100 mM sodium phosphate (pH 8.0) with 1 mM MgCl_2_. Reaction was conducted in the white opaque 96-well assay plates (Corning Costar, flat-bottom, non-treated) with a total reaction volume of 100 μl for each well. Each reaction mixture consisted of 100 nM ALDH enzyme, 0.1% DMSO, 1 mM β-mercaptoethanol, 1 mM NAD+, 1 mM acetaldehyde (lastly added) and CB29 of indicated concentration in assay buffer. Enzymatic activity was measured based on NADH-mediated fluorescence (Ex340/ Em460 nm). The activity was recorded for 10 min on a SpectraMax M2e microplate reader operated by the SoftMax Pro software. ALDH enzymatic activity caused by CB29 was normalized to the DMSO control. The single-dose inhibition data and dose-response inhibition curves were processed with the Prism software.

### LC/MS analyses of metabolites

Freshly isolated cells from murine WT and *Aldh3a1^-/-^* SMGs were used from LC/MS analyses. Please refer to **Supplemental methods** for details.

### siRNA transfection

MSGc (120,000/well) were plated in a 6 well format. 24h later, cells were transfected with 30 pmoles of siAldh3a1 (Ambion, AM16708) and scramble siRNA using Lipofectamine 3000 following Manufacturers recommended protocol. Twenty four hours later, cells were trypsinized, counted and used for downstream analyses.

### Irradiation of salivary glands and salivary collection

Thirty Gy fractionated over 5 days (6 Gy/d) were delivered to the SMG with the rest of the body lead shielded. Mice were treated with 10% Alda-341 mixed in chow or no treatment. Stimulated saliva was measured as previously described (6). Mice were anesthetized with a ketamine (80 mg/kg) and xylazine (16 mg/kg) mixture delivered by intraperitoneal injection and subcutaneously injected with 2 mg/kg pilocarpine. Saliva was collected for 15 minutes. Saliva volume was calculated by assuming the density as 1, was normalized to the mouse body weight by dividing the total collected saliva volume by the mass of the mouse (kg).

### Study approval

Human salivary glands were procured from Head and neck cancer patients in accordance to the guidelines approved by the Stanford University’s Institutional Review Board. Written informed consent was received from participants prior to inclusion in our study. The human tissue was rinsed twice with diluted betadine solution and washed with excess sterile Phosphate Buffered Saline (PBS) three times before dissociation.

### Statistics

All data are represented as averages with standard deviation (SD). Statistical ANOVA and Student’s t tests were used to compare the data. All tests performed were two-sided with an alpha level of 0.05. P ≤ 0.05 was considered to be significant. All data was analyzed using GraphPad Prism 8.4.3 (GraphPad Software Inc, La Jolla, CA).

## Acknowledgements

We would also like to thank Nan Xiao, Daria Mochly-Rosen’s group and Sarah Knox’s group for valuable input and supplies for this research work. We would like to sincerely thank Rose-Anne Romano at University of Buffalo for mSGc cells. We thank Dr. Vasilis Vasiliou for *Aldh3a1^−/−^* transgenic mice. We thank Pauline Chu at the Histology core facility at Stanford School of Medicine. We thank Dr. John Coller and Vida Shokoohi for RNA-seq. We thank Sivakumasundari V for her assistance with single cell RNA-Seq data analyses. We acknowledge John Perrino and EM core facility at Stanford University for assistance with EM data.

## Data availability statement

The datasets used and/or analyzed during the current study are available from the corresponding author on reasonable request.

## Supplementary methods

### Drugs used

Alda-341 (d-limonene) was purchased from Sigma-Aldrich (St. Louis, MO). Carbonyl cyanide 4- (trifluoromethoxy)phenylhydrazone (FCCP), 4-hydroxynonenal (4-HNE), ALDH3A1 inhibitor CB29 were purchased from MilliporeSigma (Burlington, MA) and reconstituted in sterile DMSO (FisherScientific, Hampton, NH). MitoTEMPO was purchased from MilliporeSigma and dissolved in molecular grade sterile water. Hydrazine derivative Hydralazine hydrochloride was purchased from TCI America (Portland, OR) and dissolved in molecular grade water (Thermo Fisher Scientific, Waltham, MA).

### Salivary gland isolation and culture from mouse and human SMGs

With sterile clean forceps and surgical scissors, a small cut (0.5 inch) was made vertically below the jaw line to expose the submandibular glands. Connective tissue was removed from the glands with the help of sharp forceps. The gland was minced into small pieces by using a clean surgical blade. The minced gland tissue (mouse or human) was transferred into a 15-ml tube containing 3-6 ml of dissociation media (DMEM/F12 with collagenase (0.025%) and hyaluronidase (0.04%), 6.25 mM CaCl_2_, and fungizone. Tissue dissociation was allowed for an hour at 37 C on a shaker. Equal amount of Dispase I enzyme (CORNING, Corning, NY) was added to the dissociation media and incubated for another hour. The dissociated cells were passed through a 100-micron sterile filter and were spun down at 1200 rpm for 5 minutes. After removing the supernatant, RBCs were lysed by incubating the cell suspension in ACK RBC lysis buffer (LONZA, Basel, Switzerland) for 2 minutes at room temperature. The buffer was neutralized by adding media containing FBS and cells were spun down again. To the cell pellet, 0.25% trypsin-EDTA solution was added and incubated for 1 minute at 37 C to facilitate single cell dissociation. After neutralizing with stem cell media, the cell suspension was passed through a sterile 70 micron filter. For human sample, The cells were spun down and re-suspended in required amount of salisphere media (DMEM/F12 + GlutaMax media containing 10% FBS, 1x antibiotic-antimycotic, 1% N2 supplement, 20 ng/mL epidermal growth factor-2, 20 ng/mL fibroblast growth factor-2, 10 μg/mL insulin, 1 μM dexamethasone, 10 μM Y-27632). Cell suspension after gland dissociation was mixed with twice the volume of Growth factor reduced matrigel (CORNING) and 75 μl of the mixture was added as a drop to a 12-well plate. The mixture was incubated at 37 C for 20 minutes to allow it to solidify followed by addition of 1 ml of salisphere media. Dissociated murine salivary gland cells were blocked in 3% BSA in PBS and stained for live cells (Zombie NIR dye, 1:1000) for 10 minutes on ice. Cells were washed once with cold blocking buffer and resuspended in an antibody cocktail containing anti-EpCAM-FITC, anti-CD24-APC, anti-CD31-PE and anti-CD45-PE (1:200) in blocking buffer and incubated for 10 min on ice. Cells were washed with excess blocking buffer, resuspended and subjected to FACS analyses and sorting using a BD FACS ARIAII. Unstained cell sample were used to gate the positive cells with the FACS DIVA software (BD, San Jose, CA). Cells that stained negative for zombie dye followed by negative for CD45 (immune marker), CD31 (endothelial cells) expression were selected and EpCAMhigh /CD24high cells were sorted. Matrigel was plated on bottom of a 48 well plate (CORNING) and allowed to solidify at room temperature. Cell suspension containing sorted cells in salisphere media were plated on top of the matrigel. The spheres were passaged on day 7. For passaging, matrigel containing spheres were spun down and then incubated in Dispase I solution for 45 minutes. Spheres were separated by spinning the solution down and aspirating the supernatant carefully. The spheres were dissociated by incubating them in 0.25% trypsin for 10 min followed by shearing by a 27G needle syringe. The single cells were spun down and re-suspended in sphere media ready to be plated. For quantification, brightfield images of the spheres were taking by BZ-X710 KEYENCE microscope using the z-stack function. Images were stitched together into a single image and counted by ImageJ software (NIH, Bethesda, MD).

### RNA Sequencing

RNA samples were extracted using Qiagen miRNeasy kit (QIAGEN, Hilden, Germany). The cDNA was generated from extracted RNA using the Smarter μltra Low Input RNA kit (TakaraBio, Shiga, Japan). The amplified cDNA was purified and then sheared into an avg 300 bp length using Covaris S2. Libraries were generated from the sheared cDNA using Clontech Low Input Library Prep kit. Sequencing data was generated using Illumina Hi Seq 4000 system. The processed reads were imported into BRB ArrayTools, an integrated package for the visualization and statistical analysis of gene expression data developed by Dr. Richard Simon and BRB-Array Tools Development Team (Biometric Research Program, National Cancer Institute).The log2 fold change of gene expression between the two samples were calculated using EdgeR package. MetaCore (GeneGo) was used for Gene ontology analyses of the differentially expressed genes between the two groups. For the groups WT SSPC vs (WT+Alda-341), gene counts were uploaded on Biojupies (38) to generate heatmap of differentially expressed genes and GO term classification.

### Embryo gland dissection and branching morphogenesis assay

The glands were carefully fixed in 4%PFA for 10 min followed by a wash with PBS. They were carefully embedded in 200 μl of pre-warm of Histogel (Thermo-Fisher). Once the gel solidified, it was moved to 70% ethanol and process for paraffin embedding. For branching morphogenesis assay, isolated E13.5 epithelia and mesenchyme were separated using Dispase treatment and mechanical dissection and cultured in a drop of laminin on a nucleopore filter over serum-free DMEM/F12 containing transferrin and ascorbic acid (complete media) as described for the E13.5 SMG. Epithelia were cultured with 400 ng/ml FGF10 (R&D Systems, Minneapolis, MN) and 0.5 μl/ml heparin sulfate (Sigma-Aldrich) in the presence or absence of Alda-341 or vehicle control (PEG-400) and were subjected to RNA isolation or fixed for immunostaining after 24-48 h.

### EM imaging

Samples were fixed in Karnovsky’s fixative: 2% Glutaraldehyde (EMS, Sumter, SC) and 4% paraformaldehyde (EMS) in 0.1M Sodium Cacodylate (EMS) pH 7.4 for 1 hr. The fix was replaced with cold/aqueous 1% Osmium tetroxide (EMS Cat# 19100) and were then allowed to warm to Room Temperature (RT) for 2 hrs rotating in a hood, washed 3X with μltrafiltered water, then stained in 1% Uranyl Acetate at RT 2hrs while rotating. Samples were then dehydrated in a series of ethanol washes for 30 minutes each @ RT beginning at 50%, 70% EtOH then moved to 4°C overnight. They were placed in cold 95% EtOH and allowed to warm to RT, changed to 100% 2X, then Propylene Oxide (PO) for 15 min. Samples are infiltrated with EMbed-812 resin (EMS) mixed 1:2, 1:1, and 2:1 with PO for 2 hrs each with leaving samples in 2:1 resin to PO overnight rotating at RT in the hood. The samples are then placed into EMbed-812 for 2 to 4 hours then placed into molds w/labels and fresh resin, orientated and placed into 65° C oven overnight. Sections were taken around 80nm, picked up on formvar/Carbon coated slot Cu grids, stained for 40seconds in 3.5% Uranyl Acetate in 50% Acetone followed by staining in Sato’s Lead Citrate for 2 minutes. Observed in the JEOL JEM-1400 120kV. Images were taken using a Gatan Orius 832 4k X 2.6k digital camera with 9 um pixel.

### LC/MS analyses

Cells were washed with warm media and [U-^13^C]glucose RPMI medium (10 dialyzed FBS) lacking glucose, serine, and glycine (TEKnova, Hollister, CA) and reconstituted with [U^13^C]glucose (2 g/liter), serine (0.03 g/liter), and glycine (0.01 g/liter) was added to each well and incubated for 6 hours. Cells were washed twice with ice-cold PBS prior to extraction with 80:20 acetonitrile:water over ice for 15min, sonicated for 30s Biorupter 300 (Diagenode) sonicator, then spun down at 1.5 x 104 RPM for 10 min. 200 µL of supernatant was taken out of for the LC-MS/MS analysis immediately. Quantitative LC-ESI-MS/MS analysis of ^13^C-glucose-labeled cell extracts was performed using an Agilent 1290 UHPLC system equipped with an Agilent 6545 Q-TOF mass spectrometer (Santa Clara, CA, US). A hydrophilic interaction chromatography method (HILIC) with an BEH amide column (100 x 2.1 mm i.d., 1.7 μm; Waters) was used for compound separation at 35 °C with a flow rate of 0.3ml/min. The mobile phase A consisted of 25 mM ammonium acetate and 25mM ammonium hydroxide in water and mobile phase B was acetonitrile. The gradient elution was 0 – 1.5 min, 80B; 1.5 – 7 min, 80B → 40B; 7 – 8.5 min, 40B; 8.5 – 8.7 min, 40 → 80B; 8.7 – 10 min, 80B. The overall runtime was 10 min and the injection volume was 6 μL. Agilent Q-TOF was operated in negative mode and the relevant parameters were as listed: ion spray voltage, 3500 V; nozzle voltage, 1000 V; fragmentor voltage, 125 V; drying gas flow, 11 L/min; capillary temperature, 300 °C, drying gas temperature, 320 °C; and nebulizer pressure, 40 psi. A full scan range was set at 50 to 1200 (m/z). The reference masses were 119.0363 and 980.0164. The acquisition rate was 2 spectra/s. Data processing was performed with Agilent Profinder B.08.00 (Agilent technologies). The mass tolerance was set to +/- 15 ppm and RT tolerance was +/- 0.2 min. Natural isotope abundance was corrected using Agilent Profinder software (Agilent Technologies). For normalization of ion counts, cell pellets were vacuum-dried, then protein concentration was determined by Pierce^TM^ BCA protein assay kit (Thermo-Fisher) according to manufacturer’s instructions.

### Supplementary Figure legends

**Supplemental Figure 1:**
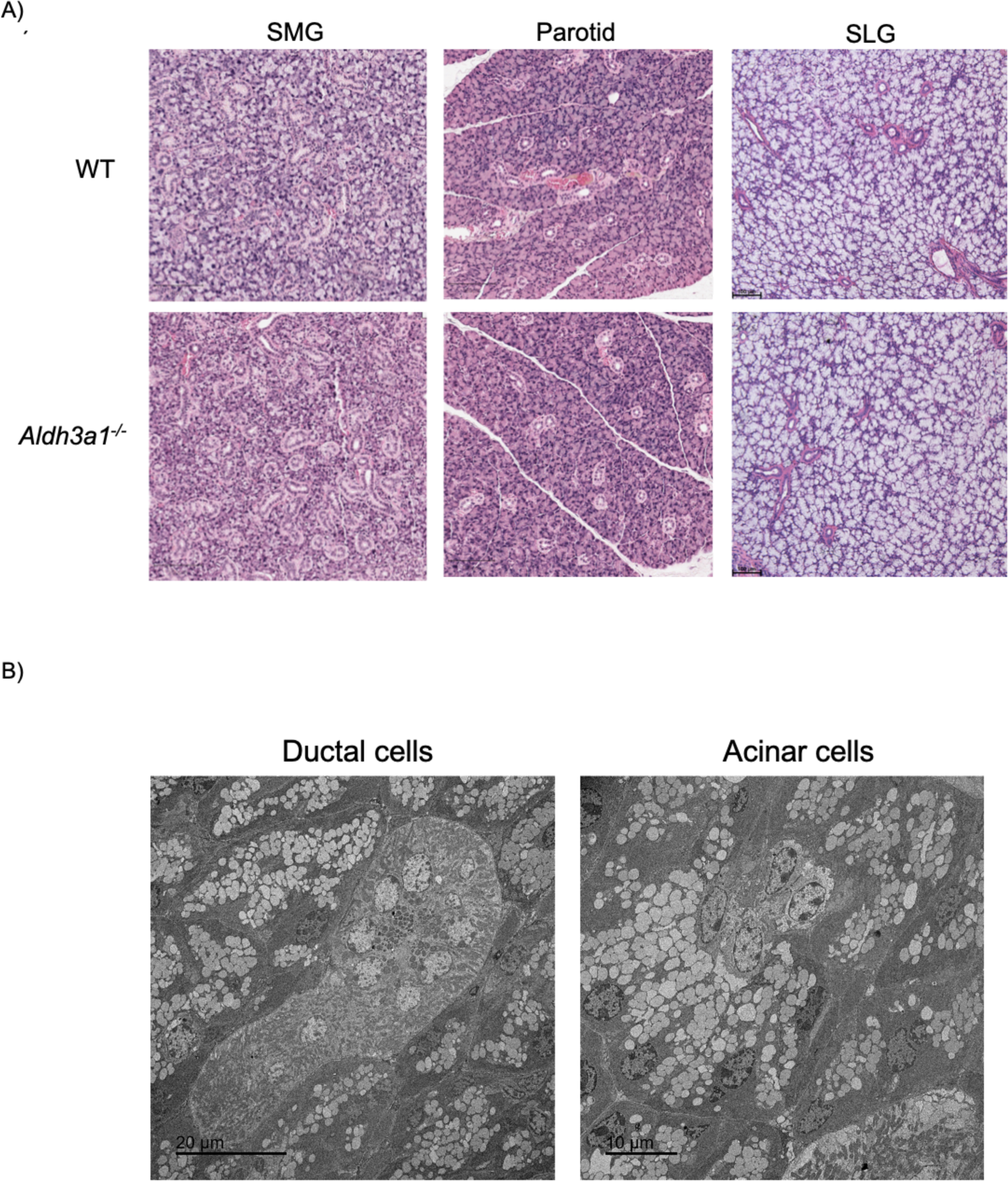
**A)** Tissue morphology of major murine salivary glands in WT and *Aldh3a1^-/-^* mice represented by hematoxylin and eosin staining imaged at 100x total magnification (n=3/group). Scale bar: 100 μM. B) Representative TEM images of ductal and acinar cells in WT SMGs (n=3).

**Supplemental Figure 2:**
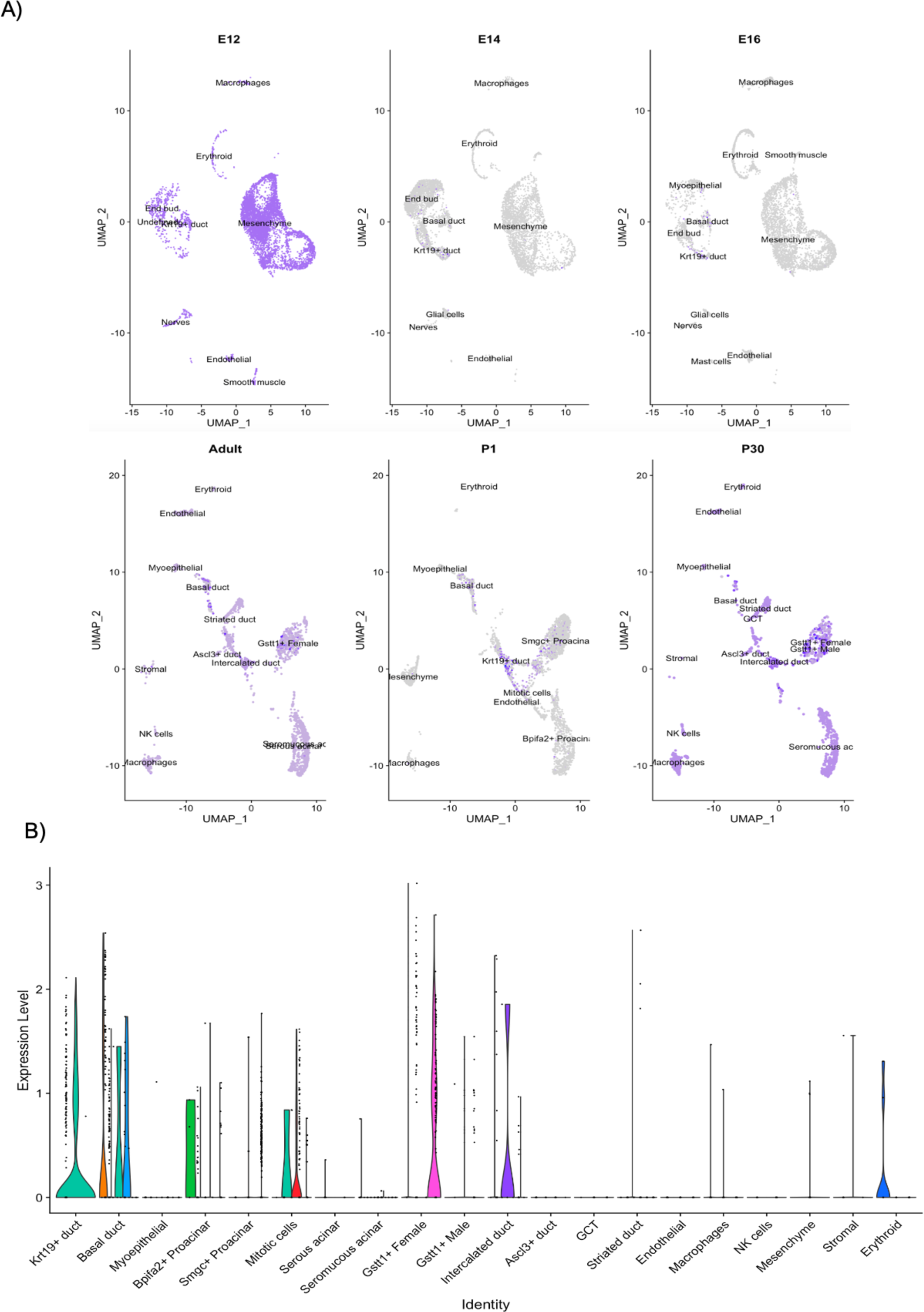
SEURAT analyses of embryonic, postnatal and adult murine SMG epithelium (38). **A)** UMAPs of Aldh3a1 expression across individual cluster of cells in embryonic (E12, E14 and E16), postnatal (P1 and P30) and adult murine SMGs. **B)** Violin plot of Aldh3a1 expression across identified cell clusters in postnatal and adult murine SMGs.

**Supplemental Figure 3:**
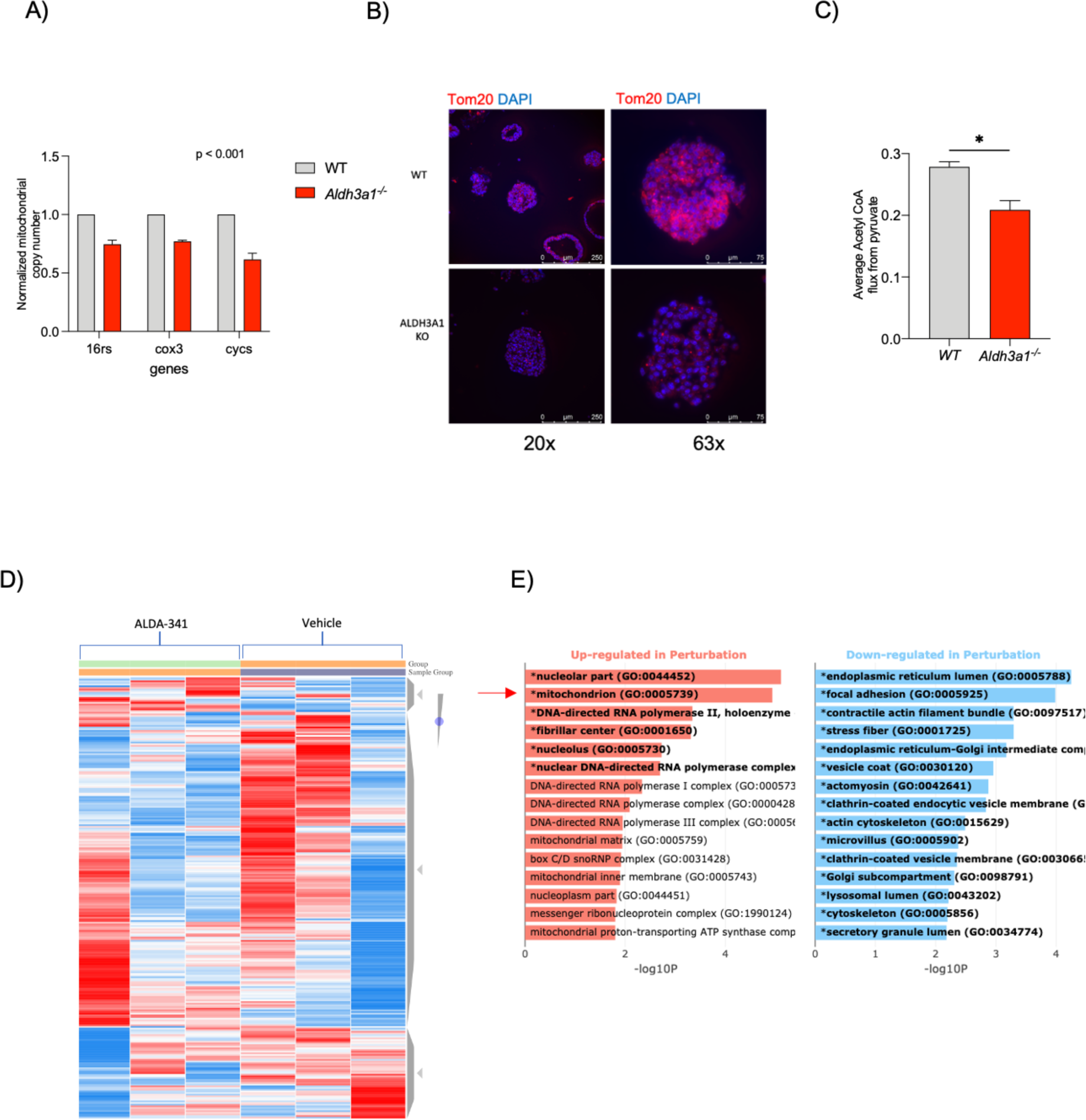
**A)** Mitochondrial DNA copy represented by DNA amount of mitochondrial encoded genes 16rs, cox3 and cycs normalized to DNA amount of nuclear encoded gene beta-globin as assessed by PCR (n=3 mice/group).**B)** Representative immunostaining images of Tom20 in WT and *Aldh3a1^-/-^* salispheres at 200x and 630x total magnification (scale bar: 250 μM for 200x image, 75 μM for 630x). **C)** Carbon labelling experiments demonstrate differences in acetyl-co A influx in WT and *Aldh3a1^-/-^* murine SSPC (n=3/group). **D)** Heatmap of differentially expressed genes in WT SSPC treated with Alda-341 and vehicle control. **E)** GO term classification of top differentially expressed genes between the two groups identified “mitochondrion” (GO:0005739) to be significantly upregulated in drug treated group as compared to vehicle control. Error bars represent SD. Student’s t-test was used to calculate the p value for panel **A** and **C.**

**Supplemental Figure 4:**
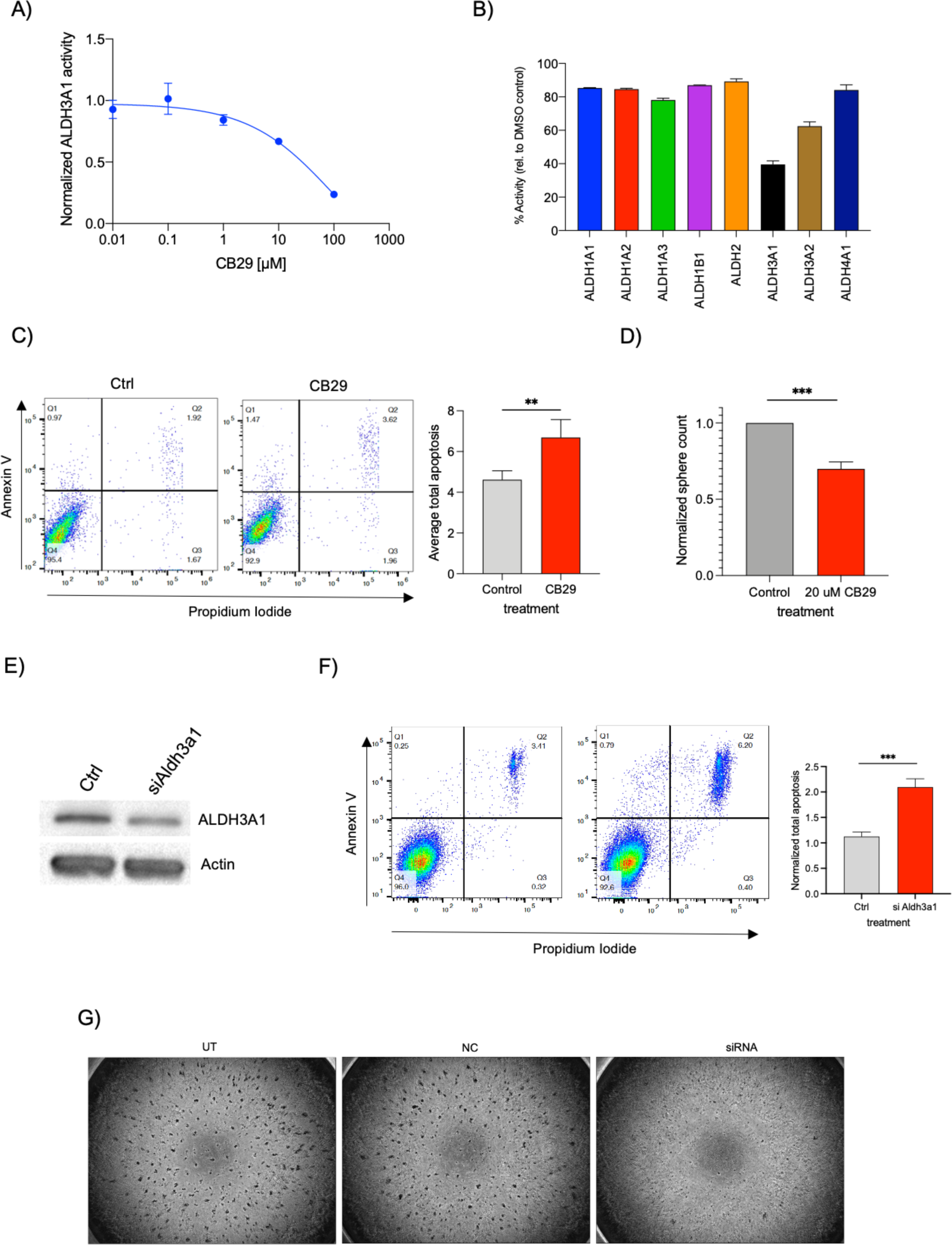
**A)** Dose response curve of CB29 on ALDH3A1 activity **B)** Specificity of CB29 (20 uM) in inhibition of ALDH3A1/2 activity demonstrated by enzyme activity assays of various ALDH isoforms (n=2, 4 technical replicates). **C)** FACS plots showing Annexin-V/PI staining of mSGc 24 hours post treatment with CB29 and vehicle control, quantified and displayed as graph in right panel (n=3, in triplicates). **D)** Average number of spheres counted at day 7 post treatment with 20 uM CB29 and vehicle control (n=2, 3 technical replicates). **E)** Western blot probed for Aldh3a1 and actin in protein isolated from mSGc treated with siAldh3a1 and control. F) FACS plots showing Annexin-V/PI staining of mSGc 24 hours post transfection with siAldh3a1 and control, quantified and shown in the right panel (n=2, in triplicates). Error bars represent SD. Student’s t-test was used to calculate the p value for panel **C, D** and **F.** Two-way ANOVA was used to determine p value for panel **B**. (** represents p value < 0.01, *** < 0.001).

**Supplementary Table 1.**

| Antibody | Source | Catalog/Clone No | Dilution | Application * |
| --- | --- | --- | --- | --- |
| Aqp5 | Alomone Labs | AQP-005 | 1:100 | IF |
| Sox9 | EMD Millipore | AB5535 | 1:100 | IF |
| Ck7 | Novus Biologicals | RCK-105 | 1:100 | IF |
| Tom20 | Santa Cruz Biotechnology | F-10 | 1:100 | IF |
| Parkin | Santa Cruz Biotechnology | PRK8 | 1:100 | IF |
| 4-HNE | R&D systems | 198960 | 1:500 | IHC, WB |
| Actin | Santa Cruz Biotechnology | C4 | 1:1000 | WB |
| EpCAM | eBioscience | G8.8 | 1:200 | FACS |
| CD24 | eBioscience | M1/69 | 1:100 | FACS |
| CD31 | eBioscience | 390 | 1:200 | FACS |
| CD45 | eBioscience | 30-F11 | 1:200 | FACS |

\* IF = Immunofluorescence
IHC = Immunohistochemistry
WB = Western Blotting
FACS = Flow cytometry analyses

